# Alternative oxidase induction protects Candida albicans from respiratory stress and promotes hyphal growth

**DOI:** 10.1101/405670

**Authors:** Lucian Duvenage, Louise A. Walker, Aleksandra Bojarczuk, Simon A. Johnston, Donna M. McCallum, Carol A. Munro, Campbell W. Gourlay

**Affiliations:** Kent Fungal Group, School of Biosciences, University of Kent, Kent, CT2 9HY, UK.; MRC Centre for Medical Mycology at the University of Aberdeen, Institute of Medical Sciences, Foresterhill, Aberdeen AB25 2ZD; Department of Infection, Immunity & Cardiovascular Disease and Bateson Centre, University of Sheffield, Firth Court, Western Bank Sheffield, S10 2TN, UK

## Abstract

The human fungal pathogen *Candida albicans* possesses two genes expressing a cyanide-insensitive Alternative Oxidase (Aox) enzymes in addition to classical and parallel electron transfer chains (ETC). In this study, we examine the role of Aox in *C*. *albicans* under conditions of respiratory stress, which may be inflicted during its interaction with the human host or co-colonising bacteria. We find that the level of Aox expression is sufficient to modulate resistance to classical ETC inhibition under respiratory stress and are linked to gene expression changes that can promote both survival and pathogenicity. For example we demonstrate that Aox function is important for the regulation of filamentation in *C*. *albicans* and observe that cells lacking Aox function lose virulence in a zebrafish infection model. Our investigations also identify that pyocyanin, a phenazine produced by the co-colonising bacterium *Pseudomonas aeruginosa*, inhibits Aox-based respiration in *C*. *albicans*. These results suggest that Aox plays important roles within respiratory stress response pathways which *C*. *albicans* may utilise both as a commensal organism and as a pathogen.

## Introduction

*C. albicans* is a commensal yeast which is found in the majority of the population. However it is also a major fungal pathogen of humans and a leading cause of hospital-acquired infections which have a high mortality rate (Edmond et al., 1999). *C*. *albicans* is Crabtree Negative yeast that relies largely on respiration to produce energy for growth and is highly adaptable in its use and assimilation of a variety of carbon sources (Ene et al., 2012). Recent work has identified a number of fungal-specific subunits of mitochondrial Complex I raising the possibility of the development of novel antifungals that target this organelle (She et al., 2015). Although the importance of mitochondrial function for growth and proliferation of *C*. *albicans* is recognised, the influence of alternative pathways of respiration remain to be fully explored.

In addition to the classical electron transfer chain (ETC), *C*. *albicans* possesses a cyanide-insensitive alternative pathway, not found in mammals, which permits respiration when the ETC is inhibited (Huh and Kang, 1999). This pathway relies on an alternative oxidase (Aox), of which two isoforms exist. Aox1 is a low-abundance isoform that is constitutively expressed while Aox2 can be rapidly induced in response to inhibition of classical respiration (Huh and Kang, 2001). Aox activity is not coupled to the generation of a proton gradient across the mitochondrial membrane, and thus alternate respiration produces significantly less ATP than the classical oxidative phosphorylation (Helmerhorst et al., 2002). This suggests that Aox based respiration does not play a key role in energy production but likely permits respiration under conditions of classical chain inhibition. Metabolomic analyses indicate that inhibition of the ETC is associated with changes in the purine nucleotide cycle, as well as a depletion in TCA cycle intermediates and aspartate (Balcke et al., 2011). In this context Aox induction may be crucial as respiration provides electron acceptors to support the biosynthesis of aspartate, which is considered as one of the limiting factors for cell proliferation (Sullivan et al., 2015). Sustained respiration is also important as mitochondria play a key role in the synthesis of several lipids that support growth and cell maintenance. The import of phosphatidylserine into mitochondria to generate phosphatidylethanolamine, an important constituent of the plasma membrane, is an example of this function (Miyata et al., 2016). The ability of Aox to maintain respiration under conditions of classical ETC inhibition is therefore likely to support biosynthetic pathways required for growth.

An alternative oxidase system is found in all plants, most fungi, algae, and some protists (Rogov and Zvyagilskaya, 2015). Aox is located in the inner mitochondrial membrane where it receives electrons from reduced ubiquinone and catalyzes the reduction of oxygen to water. The catalytic core of Aox is formed by a four-α-helix bundle, containing the di-iron catalytic site, flanked by two additional α-helices anchoring the protein to the membrane (Siedow and Umbach, 1995). Studies of *Pichia stipitis* and *Neurospora crassa* indicate that fungal Aox differs from Aox in plants in that it occurs as a monomer and is not induced by α-keto acids such as pyruvate (Umbach and Siedow, 2000). Another unique feature of fungal Aox is that its activity is stimulated by purine nucleotides such as AMP, ADP and GMP (Sakajo et al., 1997; Vanderleyden et al., 1980). The regulatory differences observed may suggest that Aox participates within different physiological processes in fungal and plant species.

In plant fungal pathogens such as *Ustilago maydis, Moniliophthora perniciosa* and *Sclerotina sclerotiorum*, Aox is reported to be more active during the mycelial growth phase suggesting that the metabolic control afforded by alternative respiration is an important factor in morphogenesis (Juárez et al., 2006; Thomazella et al., 2012; Xu et al., 2012). In the human fungal pathogen *Aspergillus fumigatus* AoxA was found to contribute to oxidative stress resistance and to resistance to killing by macrophages (Magnani et al., 2008) but was not essential for virulence *in vivo* (Grahl et al., 2012). In *Cryptococcus neoformans* the sole alternative oxidase Aox1 is induced at 37 °C and plays a significant role in virulence in the murine inhalation model (Akhter et al., 2003). In *Paracoccidioides brasiliensis* Aox has roles in morphological transition and in virulence by mediating resistance to oxidative stress imposed by immune cells (Hernández et al., 2015; Hernández Ruiz et al., 2011).

It has been proposed that in *C*. *albicans* Aox may also function in oxidative stress resistance, as oxidative agents can induce *AOX2* expression (Huh and Kang, 2001). This is supported by the observation that the induction of Aox2 provides resistance to fluconazole, which has been shown to elevate ROS levels as part of its fungistatic action (Yan et al., 2009). Inhibition of Aox with benzhydroxamic acid significantly raised the ratio of oxidised/reduced glutathione, suggesting that Aox may also have a role in maintaining redox balance (Ruy et al., 2006). Controlling the partitioning of electrons between cytochrome c oxidase and Aox has also been proposed to provide the respiratory system with flexibility regarding ATP demand and maintaining cellular redox balance (Vanlerberghe, 2013). Although the contribution of the alternative pathway to total respiration is minimal in dividing yeast cells, it has been reported to be more active in hyphal (Guedouari et al., 2014) and aged cells (Helmerhorst et al., 2005).

Given that Aox is not found within mammals it has been proposed as a potential drug target for control of plant fungal pathogens. However Aox support of respiration is reported as one of the mechanisms of resistance to QoI antifungals (which inhibit fungal respiration by binding to the Qo site of the cytochrome *bc1* complex of the classical ETC) proposed for use to control plant pathogens (e.g. the strobilurins) (Fernández-Ortuño et al., 2008). Examples of plant fungal pathogens in which alternative respiration was shown to confer resistance include *Zymoseptoria tritici* (Miguez et al., 2004) and *Botrytis cinerea* (Ishii et al., 2009). Aox has also been targeted in the eukaryotic parasite *Trypanosoma brucei*, which cause African sleeping sickness (Menzies et al., 2018). Therapies targeting Aox in these organisms using drugs such as salicylhdroxamic acid (SHAM) or ascofuranone have demonstrated efficacy of Aox inhibition with small molecules (Brohn and Clarkson Jr., 1978; Yabu et al., 2003). The hydroxynaphthoquinone class of molecules, including Atovaquone, also act by inhibiting the Qo site within the ETC and have been shown to be effective against malaria parasites as well as the opportunistic fungal pathogen *Pneumocystis jirovecii* (Fisher and Meunier, 2008). Alternative oxidase inhbitors were shown to be synergistic with Atovaquone against *Plasmodium falciparum* (Murphy and Lang-unnasch, 1999), showing that simultaneous inhibition of the classical and alterative pathways is more effective, and highlights the importance of Aox as a drug target.

The clear necessity of the alternative pathway for resistance to inhibition of classical respiration may be relevant to *C*. *albicans* fitness and survival under certain conditions. For example, nitric oxide (NO) is produced by phagocytes as a broad-spectrum defence against microbes. NO and reactive nitrogen species such as peroxynitrite produced by macrophages in response to *C*. *albicans* can inhibit the classical respiratory pathway (Sharpe and Cooper, 1998). As Aox is not sensitive to NO it’s expression may promote viability and escape from phagocytic cells. Certain bacteria that are found to colonise similar niches as fungal pathogens are known to produce compounds which inhibit respiration. An example is cyanide production by *Pseudomonas aeruginosa* in the cystic fibrosis lung (Lenney and Gilchrist, 2011), in which *C*. *albicans* is a common co-isolate (Chotirmall et al., 2010). Alternative respiration may therefore also function as a defence against respiration inhibition by competing microorganisms within mixed microbial communities.

In this study we examine the role of Aox in the resistance of *C*. *albicans* to respiratory stress which it may encounter within the human host and in co-existence with other microbes. We find that Aox activity is rapidly induced following exposure to respiratory stress, is important for hyphal induction and virulence and is targeted by the pyocyanin, a product of *P*. *aeruginosa*. Our data suggests Aox function may be important for both the commensal and pathogenic properties of *C*. *albicans* cells, providing a plausible explanation for its conservation.

## Materials and Methods

### Growth conditions and chemicals

*C*. *albicans* strains were maintained on YPD agar plates and grown in YPD in a 30 °C shaking incubator unless stated otherwise. The concentrations of sodium nitroprusside dihydrate (SNP) and salicylhydroxamic acid (SHAM) (Cat. No.’s 1.06541 and S607, Sigma-Aldrich, Dorset, UK) used for inhibition were 1 mM and 0.5 mM respectively, and were added to log phase cells followed by 18 h growth unless stated otherwise. Potassium cyanide, Calcofluor White, Congo Red, pyocyanin (Cat. No. P0046) and methylene blue were obtained from Sigma-Aldrich.

### Construction of AOX deletion, re-integration and overexpression strains

The strains used in this study are summarised in Table 1. The *aox1-aox2*Δ mutant was constructed from the wild type strain SN87 using the strategy as described in (Noble and Johnson, 2005). Briefly, *LEU2* and *HIS1* were amplified from plasmids pSN40 and pSN52 respectively, using universal primers with 80 bases homologous to the 5’ end of *AOX2* and the 3’ end of *AOX1* ORFs. This strategy was designed to delete both *AOX2* and *AOX1* simultaneously as well as the region between these two adjacent genes. The primers used were AOX2-UP2 and AOX1-UP5 (Table 2). *AOX2* only was disrupted to create the *aox2*Δ using a similar strategy by homologous recombination at the 5’ and 3’ ends of the *AOX2* open reading frame, using the primers AOX2-UP2 and AOX2-UP5. The PCR products were then transformed sequentially into *C*. *albicans* SN87 using an electroporation protocol (Thompson et al., 1998), followed by selection on agar plates containing Yeast Nitrogen Base without amino acids supplemented with 2% glucose and -His or -Leu dropout powder (Formedium, UK) as appropriate.

*AOX2* ORF was amplified from genomic DNA using *AOX2* ORF F and *AOX2* ORF R (Table 2). The PCR product was digested with BamI and XhoI (Promega, Madison, WI) and cloned into pNIM-1 (Park et al., 2005) replacing GFP. This construct was transformed into *C*. *albicans* strains by integration at the *ADH1* locus as described in (Park et al., 2005). Cells were grown at 30 °C in YPD for 16 h supplemented with 50 µg/ml doxycycline to induce expression of *AOX2*. To verify the disruption or reintegration of *AOX2*, genomic DNA was extracted from *C*. *albicans* strains using standard the phenol-chloroform method (insert reference). The *AOX2* gene was amplified using AOX2 ORF F and AOX2 ORF R.

To examine induction of *AOX2* by KCN, *C*. *albicans* strains were grown to log phase in YPD and then treated with 1 mM KCN. Samples were taken at specified time points and RNA was extracted using a E.Z.N.A.® Yeast RNA Kit (Omega Bio-tek, Norcross, GA) using the manufacturer’s instructions. cDNA was prepared using a Bio-Line SensiFAST™ cDNA Synthesis Kit (Cat. No. BIO-65053) as per the manufacturer’s instructions. Standard PCR was then performed with 2 ng template cDNA per reaction, using AOX2 ORF F and AOX2 ORF R.

### RNA isolation and RNA sequencing

*C*. *albicans* wild-type SC5314 and *aox2*Δ were grown for 5 hours in YPD at 30 °C in a shaking incubator. SHAM or Pyocyanin was added to wild-type cells to a final concentration of 0.5 mM or 80µM respectively and cells were returned to the incubator for a further 30 min. RNA was then extracted using an E.Z.N.A.® Yeast RNA Kit using the manufacturer’s instructions, for two biological replicates per group. RNA was sent to the Centre for Genome Enabled Biology and Medicine (Aberdeen, UK), which performed preparation of stranded TruSeq mRNA libraries, QC/quantification and equimolar pooling, and sequencing on an Illumina NextSeq500 with 1×75bp single reads and average depth of 30M reads per sample. Raw RNAseq data was analysed using the suite of tools available in the Galaxy platform (Afgan et al., 2016). Briefly, reads were aligned to Assembly 21 of the *C*. *albicans* genome (Candida Genome Database (Skrzypek MS, Binkley J, Binkley G, Miyasato SR, Simison M)) using HISAT2. Differentially expressed genes between untreated and treated samples were identified using Cuffdiff v2.1.1 (Schirmer et al., 2011). The p-values generated by Cuffdiff’s statistical algorithm were adjusted using Benjamini-Hochberg correction for multiple-testing to generate the q-value (allowed false discovery rate of 0.05). A q-value less than 0.05 was considered statistically significant. The data discussed in this publication have been deposited in NCBI’s Gene Expression Omnibus and are accessible through GEO Series accession number GSE117717.

### High Resolution whole cell respirometry

Respirometry was carried out in real time using an Oxygraph-2k respirometer (Oroboros Instruments, Austria) which was calibrated at 30 °C as per the manufacturer’s instructions. Cells from an overnight culture grown in YPD were added to 3 ml fresh YPD to a final optical density at 600 nm (OD_600_) of 0.2. The cells were incubated at 30 °C with shaking for 2 h. The cells were then counted and diluted in YPD to give a final cell concentration of 1 × 10^6^ cells/ml, of which 2.5 ml was added to each chamber in the respirometer. The respiration was allowed to reach routine levels before the addition of SNP and SHAM, at which time the oxygen concentration was typically 150 ppm. Routine respiration was measured immediately prior to the addition of inhibitors. The recovery of respiration was measured 20 minutes after the first addition of inhibitors. SNP and SHAM were added sequentially to 1- and 2 mM final concentrations. Pyocyanin was added to a final concentration of 80 µM. Potassium cyanide was added where indicated to give a final concentration of 2 mM. Data was analysed using Datlab 6 software (Oroboros Instruments). Student’s t-test was used to compare groups.

### Western blotting

Cells from an overnight culture in YPD were diluted in YPD to a final OD_600_ of 0.2. The cells were grown for 5 hours at 30 °C with shaking. One millimolar SNP, 1 mM KCN or 1 mM SHAM were added and samples were taken after 2 h. To examine the timescale of Aox2 expression in response to 1 mM SNP, samples were taken at 10, 20, 30 and 120 minutes to obtain cell pellets of 30 mg fresh weight. Total protein was extracted by homogenisation at 4 °C with glass beads in the presence of 125 mM Tris-HCl, 2% SDS, 2 % glycerol, 0.14 M 2-mercaptoethanol, bromophenol blue buffer at 4 °C. The samples were run on a 5% stacking, 12.5% resolving SDS-polyacrylamide gel. Proteins were transferred to PVDF membrane using a semi-dry transfer system (Bio-Rad, Watford, UK). A monoclonal antibody against *Sauromatum guttatum* Aox which recognises *C*. *albicans* Aox2 was used for Aox immunoblotting (1:100) (AS10 699, Agrisera, Sweden). Secondary binding of Anti-mouse-HRP antibodies (1:5000) (Sigma-Aldrich A9917) was detected by ECL and images captured using a Syngene GBox Chemi XX6 system.

### Hyphal growth assays

Cells from *C*. *albicans* SC5314, *aox2-aox1*Δ and *aox2-aox1Δ::AOX2* cultures grown for 16 h in YPD supplemented with 50 ug/ml doxycycline were washed three times in PBS and used to inoculate RPMI-1640 medium (Sigma-Aldrich, R8755) to a final OD_600_ of 0.1. The cells were incubated in a 180 rpm shaking incubator 37 °C for 16 h. Calcofluor White (10 µg/ml) was added to aid in visualisation and cells were examined by microscopy using a DAPI filter. The percentage of cells with germ tubes was manually determined using ImageJ v1.50 (NIH, Bethesda, MD). A total of least 500 cells were counted for each experiment.

Hyphal induction in serum-containing medium was carried out as follows: *C*. *albicans* wild-type or *AOX2* overexpression cultures grown for 16 h in YPD supplemented with 50 ug/ml doxycycline were washed three times in PBS and the OD_600_ was measured. Cells were added to DMEM + 10 % FBS (Cat. No.’s 10569010 and 10082147, Gibco, Thermo Fisher Scientific) with 10 µg/ml Calcofluor White to a final OD_600_ of 0.1 and added to a µ-Slide 8 Well (0.3 ml per well). Following 90 minutes incubation at 37 °C, 5% CO_2_, the cells were examined by microscopy using a DAPI filter. The percentage of cells with germ tubes was determined manually using ImageJ. A total of least 500 cells were counted for each experiment. Student’s t-test was used to compare groups.

### Cell wall perturbing agents susceptibility assay

To assess differences in susceptibilities to cell wall perturbing agents between the wild-type and *aox2-aox1*Δ, YPD plates were prepared containing 25 µg/ml Calcofluor White or 50 µg/ml Congo Red. YP-glycerol plates were used to assess sensitivities to 10 µg/ml methylene blue and 1 mM SNP. Cells from an overnight culture in YPD were washed three times in PBS and diluted in PBS to a final OD_600_ of 0.2. Cells were serially diluted (1:10 dilutions) and equal volumes were spotted on the plates using a replica plating tool. The plates were incubated for 48 h at 30 °C and photographed. Dectin-1 staining of cells and microscopy was carried out as described in (Duvenage et al., 2018).

### Phagocytosis assay

J774.1 murine macrophages were maintained in DMEM with 10% FBS, 200 U/ml penicillin/streptomycin (respectively, Cat No. 10569010, 10082147, 15070063, Gibco, Thermo Fisher) at 37 °C, humidified 5% CO_2_. Cells were counted and diluted in fresh medium to give 5 × 10^4^ cells in 0.3 ml which was added to the wells of a µ-Slide 8 Well (ibidi GmbH, Germany). The cells were then incubated overnight. For SHAM pre-treatment of *C*. *albicans*, 0.5 mM SHAM was added to log-phase cultures in YPD and grown for a further 18 h. Cells from an overnight *C*. *albicans* culture were washed three times in PBS and counted, then diluted to 1.5 × 10^4^ cells in 0.3 ml in macrophage culture medium with 10 µg/ml Calcofluor White and vortexed briefly. This cell suspension was then added to macrophages. The cells were then co-incubated for 1h and examined by microscopy. Uptake of *C*. *albicans* was manually assessed using ImageJ. The percentage uptake was determined as the number of internalised *C*. *albicans* relative to the total number of *C*. *albicans* cells. At least 200 *C*. *albicans* cells were counted for each experiment. Student’s t-test was used to compare groups.

### Zebrafish larva survival assay

*C*. *albicans* strains were grown overnight in YPD at 30 °C and then washed three times in PBS, counted and adjusted to give 100 or 500 colony forming units (CFU) in 1 nl and pelleted by centrifugation. Pellets were resuspended in 10% Polyvinylpyrrolidinone (PVP), 0.5% Phenol Red in PBS. Wild-type zebrafish larvae (*Nacre*) were injected with a total of 100 or 500 CFU using the method described in (Bojarczuk et al., 2016). Injected animals were maintained in E3 in 96 well plates at 28 °C. Survival was assessed by inspection of heartbeat and motility after 24 and 48 hours. Survival curves were analysed by Log-rank (Mantel-Cox) test using GraphPad Prism 7.03. Three independent experiments were performed each using 10 zebrafish larvae per group.

### Murine infection model and assessment of virulence

The virulence of *aox2-*aox1Δ was examined using a murine intravenous challenge assay. Wild-type (SC5314) and *aox2-aox1*Δ *C*. *albicans* were incubated overnight in YPD at 30 °C with shaking. BALB/c female mice (6-8 weeks old, Envigo UK) were randomly assigned into groups of 6, with group size determined by power analyses using data previously obtained using this infection model. Mice were acclimatized for 5 days prior to the experiment. Mice were weighed and tail-marked using a surgical marker pen for identification. Food and water was provided *ad libitum*. *C*. *albicans* cells were washed twice with sterile saline and diluted in sterile saline to produce an inoculum of 4 ×10^4^ CFU/g mouse body weight in 100 µl PBS. Inoculum level was confirmed by viable plate count on Sabouraud Dextrose agar. Mice were weighed and checked daily until day 3 post-infection when all mice were culled by cervical dislocation. The kidneys were removed aseptically for fungal burden determination, with kidneys weighed and homogenised in 0.5 ml sterile saline. Dilutions were plated on Sabouraud Dextrose agar and incubated overnight at 35 °C. Colonies were counted and expressed as colony forming units (CFU) per g of kidney. Change in weight of individual mice was calculated as percentage weight change relative to starting weight. An outcome score was calculated based upon kidney burdens and weight loss at time of culling (MacCallum et al., 2010). Student’s t-test was used to compare groups.

### Ethics statement

Mouse experiments were carried out under licence PPL70/9027 awarded by the UK Home Office to Dr Donna MacCallum at the University of Aberdeen. All experiments conform to the UK Animals (Scientific Procedures) Act (ASPA) 1986 and EU Directive 2010/63/EU. Zebrafish work was performed following UK law: Animal (Scientific Procedures) Act 1986, under Project License PPL 40/3574 and P1A4A7A5E. Ethical approval was granted by the University of Sheffield Local Ethical Review Panel.

## Results

### AOX is rapidly induced upon ETC inhibition to support respiratory function

Our aim was to investigate the regulation and physiological roles of AOX in *C*. *albicans*. To facilitate this we adopted a double deletion strategy to remove both *AOX* genes (AOX1 and *AOX2*) and produce a strain devoid of alternative oxidase activity. The deletion of both *AOX* genes was confirmed by PCR (Supplementary Material Figure S1A, B) and RT-PCR (Supplementary Material Figure S1C). Next we sought to identify conditions that could be used to perform experiments under conditions of AOX inhibition; *AOX2* could be rapidly induced upon application of COX complex inhibitors, with an increase in *AOX2* mRNA detectable within 15 min of KCN exposure (Figure 1A). Aox2 was strongly induced by the Nitric Oxide donor Sodium Nitroprusside (SNP) and to a similar degree upon cyanide treatment (KCN) treatment which both target the ETC (Figure 1B). The alternative oxidase inhibitor SHAM, used in later experiments, did not induce Aox2 expression (Figure 1B). Aox2 protein was detectable within 20 min of induction and increased in level over time (Figure 1C). Aox2 was not detectable in untreated cells by western blot.

**Figure 1.**
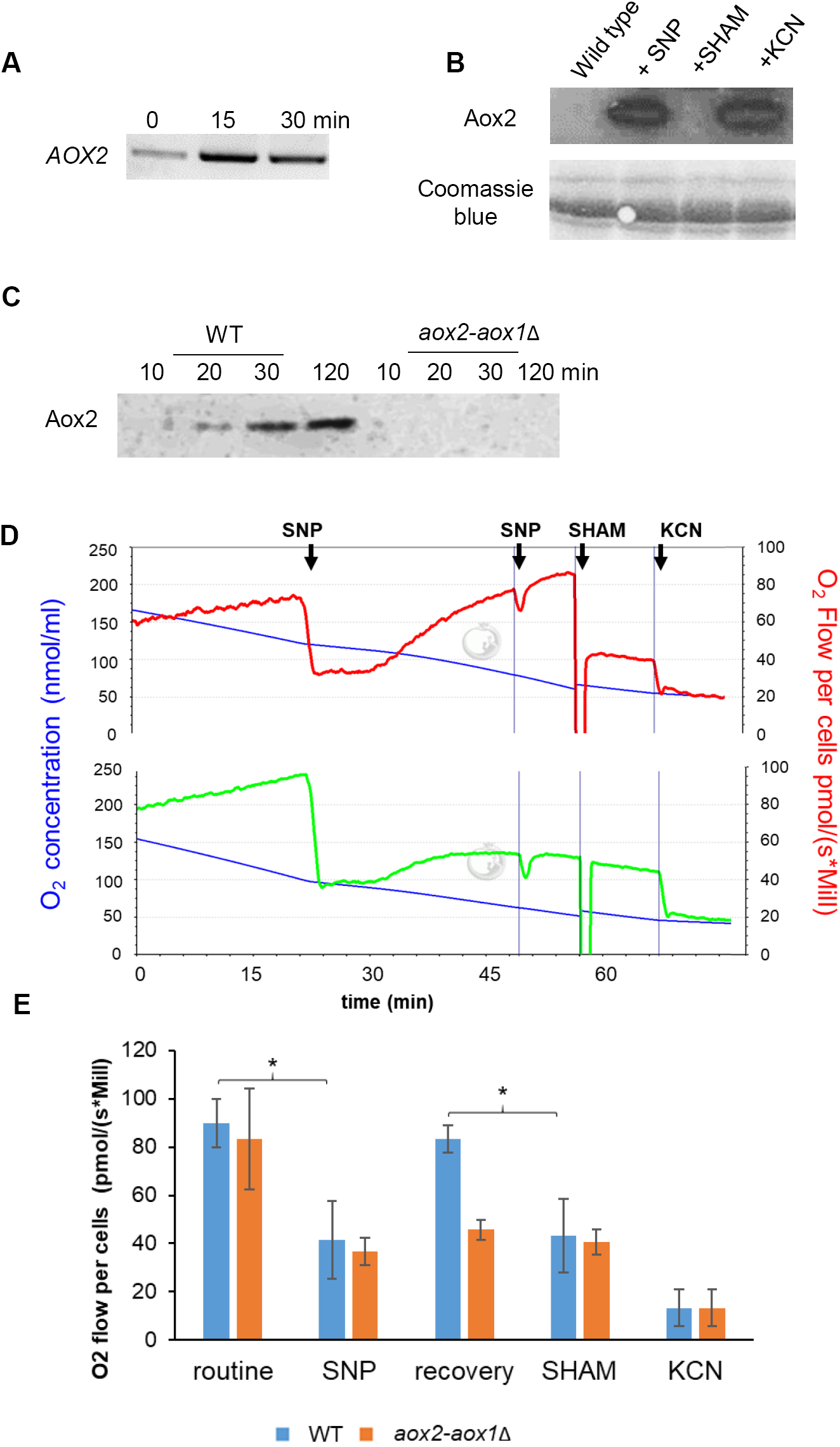
AOX restores respiration rapidly following ETC inhibition. **A**. RT-PCR was used to monitor *AOX2* expression after 15 and 30 min exposure to 1 mM KCN**. B.** Immunoblot of Aox2 following 1 mM KCN, 1 mM salicylhydroxamic acid (SHAM) or 1 mM sodium nitroprusside (SNP) treatment for 2 h. Coomassie stained gel is shown as a loading control. **C.** Immunoblot to monitor Aox2 protein over time following SNP exposure in the wild-type strain and the *aox2-aox1*Δ mutant. **D.** Respirometry analysis of the wild-type strain and *aox-aox1*Δ mutant. Routine respiration was recorded immediately prior to the addition of drugs. SNP and SHAM were added to give 1- and 2 mM final concentrations. A final addition of 2 mM KCN was used to inhibit remaining respiration. Respiration measurements were recorded 5 min after addition of each respective drug. The recovery of respiration was measured 20 min after the first addition of SNP. **E.** Summary of data from three independent respirometry experiments. Results are presented as means ± standard deviation. Student’s t-test was used to compare groups, *p<0.01.

In order to study the physiological effects of Aox induction whole-cell respirometry was performed using wild-type and *aox2-aox1*Δ strains. While wild type cells recovered full respiratory function within approximately 25 min following SNP treatment, cells lacking Aox function failed to do so (Figure 1D). Respiration observed in wild type cells that had recovered following SNP treatment was insensitive to further SNP addition but could be fully inhibited by the Aox inhibitor SHAM (Figure 1D and 1E). The *aox2-aox1*Δ mutant was insensitive to further addition of SNP following initial treatment with this inhibitor and was also unaffected by SHAM addition following SNP treatment (Figure 1D and 1E). Similar results were observed when KCN was used instead of SNP (Supplementary Figure S2). These data are in line with the observed upregulation of Aox mRNA and protein following ETC inhibitor application and confirm that the subsequent restoration of respiration is Aox dependent. The importance of Aox depedent respiration under ETC inhibition is highlighted by our finding that *aox2-aox1*Δ cells fail to grow on agar plates containing SNP or methylene blue (Supplementary Figure S1D), a commonly used antifungal agent that uncouples classical respiration by transferring electrons from the ubiquinone pool directly to oxygen (Schirmer et al., 2011).

To allow for the examination of the effects of AOX induction we introduced *AOX2* into both a wild-type and *aox2-aox1*Δ background under the control of an inducible promoter. This approach enabled us to examine the effects of the constitutive overexpression of Aox2, or Aox2 overexpression in addition to endogenous Aox2 that could be induced by classical ETC inhibition. Overexpression of Aox2 in the *aox2-aox1*Δ background reduced the magnitude of respiration inhibition by KCN and the remaining respiration was sensitive to SHAM, confirming Aox2 activity (Figure 2A and 2B). When Aox2 was overexpressed in the wild-type background, addition of cyanide did not cause a decrease in respiration (Figure 2C). The decrease in respiration level following SHAM addition showed that the majority of the respiration was due to alternative oxidase activity (Figure 2C). These data suggest that the Aox levels correlate well with protection against agents that inhibit the ETC.

**Figure 2.**
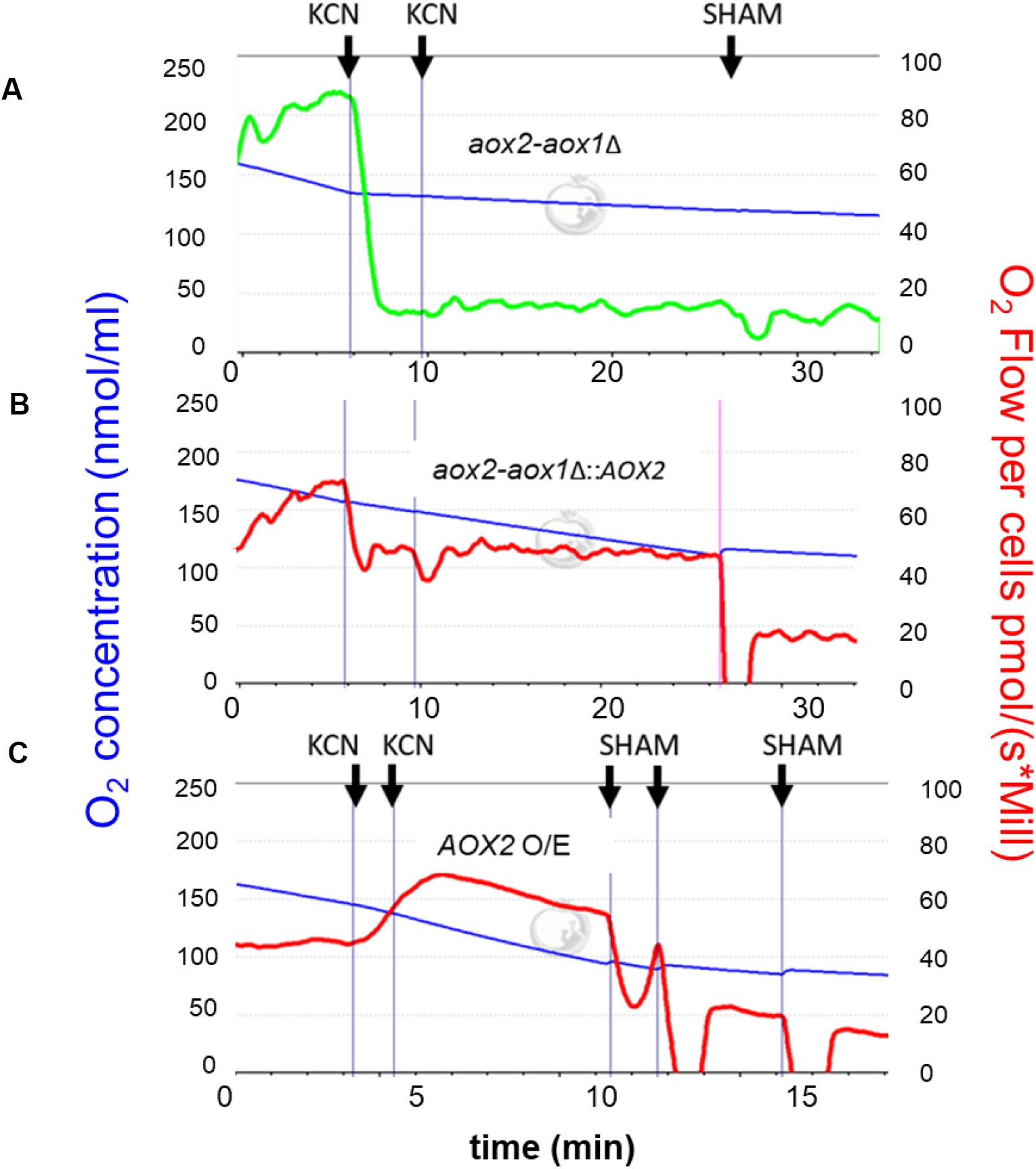
Increased *Aox* activity elevates resistance to ETC inhibition. Representative examples of respirometry experiments using *AOX2* overexpression strains: **A.** *aox2aox1*Δ control, **B.** *aox2aox1*Δ::*AOX2*, **C.** *AOX2* overexpression strain. KCN and SHAM were added sequentially to 1- and 2 mM final concentrations. The blue line shows oxygen concentration and red/green lines show the O_2_ -flux.

### The absence of Aox does not affect cell wall integrity or macrophage recognition

Several studies have shown a link between respiration and cell wall integrity in *C*. *albicans* (Dagley et al., 2011; Khamooshi et al., 2014; She et al., 2013). To determine whether alternative respiration contributes to cell wall integrity, we examined the sensitivity of *aox-aox1*Δ to the cell wall damaging agents Calcofluor White and Congo Red. The growth of *aox2-aox1*Δ cells was similar to the wild-type upon exposure to these agents (Figure 3A). We also examined surface exposure of chitin and β-glucan by cell wall staining with wheat-germ agglutinin and dectin-1 but found no significant difference between wild type and *aox2-aox1*Δ cells (Supplementary Figure S3).

**Figure 3.**
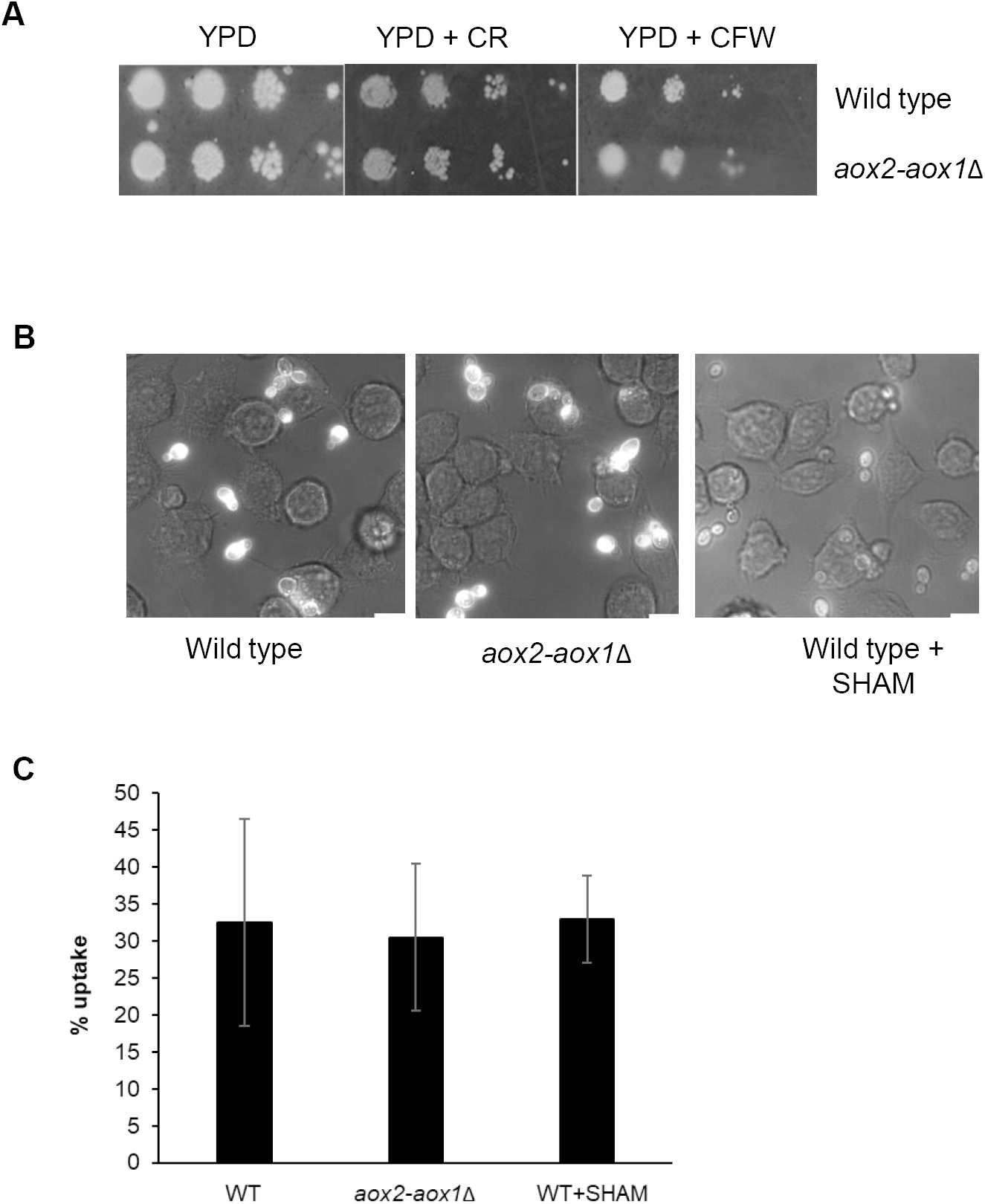
**Loss of AOX function does not affect the cell wall integrity or macrophage recognition A.** The wild-type or *aox-aox1*Δ mutant were spotted onto plates containing 25 µg/ml Calcofluor White (CFW) or 50 µg/ml Congo Red. **B. W**ild-type *C*. *albicans* were grown with or without treatment with 1 mM SHAM for 18 h, washed and co-incubated with J774.1 macrophages. Representative examples of uptake by macrophages are shown. **C.** Summary of *C*. *albicans* uptake by macrophages for three independent experiments. At least 200 *C*. *albicans* cells were counted per experiment. Results are presented as means ± standard deviation. Student’s t-test was used to compare groups.

A difference in recognition and uptake by macrophages may also reflect cell wall changes as this relies on recognition of surface exposed cell wall PAMPS, especially β-glucan (Gow et al., 2007). We examined the uptake of *aox2-aox1*Δ and the wild-type strain that had been grown in the presence of the Aox inhibitor SHAM by murine macrophages. The percentage uptake by macrophages of *aox2-aox1*Δ or SHAM-treated wild-type cells was not significantly different to that of the untreated wild-type strain (Figure 3B and 3C). Overall these data suggest that suppression of alternative respiration alone, either by chemical means or by deletion of the alternative oxidases, does not affect cell wall integrity or *C*. *albicans* immune cell recognition.

### Alternative respiration has a role *C*. *albicans* hyphal transition

Mitochondrial activity has been shown to modulate Ras1-Cyr1-PKA pathway (Grahl et al., 2015) and so may influence yeast-to-hyphae transition in *C*. *albicans*. In addition, loss of Complex I activity of the ETC has been linked to a reduction in the ability to activate the hyphal transition program in *C*. *albicans* (McDonough et al., 2002). Mitochondrial function is therefore clearly linked to the hyphal transition programme and we sought to determine whether Aox plays a role in this process. Under conditions known to activate the hyphal transition programme we found that *aox2-aox1*Δ mutant cells displayed a defect in germination. After 16 h growth in RPMI at 37 °C significantly fewer *aox2-aox1*Δ cells had germinated when compared to the wild type strain, and a high proportion of *aox2-aox1*Δ yeast cells exhibited an aberrant elongated morphology (Figure 4A, B). This effect could be reversed by re-expression of *AOX2* (Figure 4B). Further evidence for a direct role of Aox in activating hyphal transition was obtained by examining the effects of overexpression of *AOX2*. Overexpression of *AOX2* in DMEM + 10% serum caused a significant increase in the proportion of cells which had formed germ tubes after 90 min when compared to the control (Figure 4C, D).

**Figure 4.**
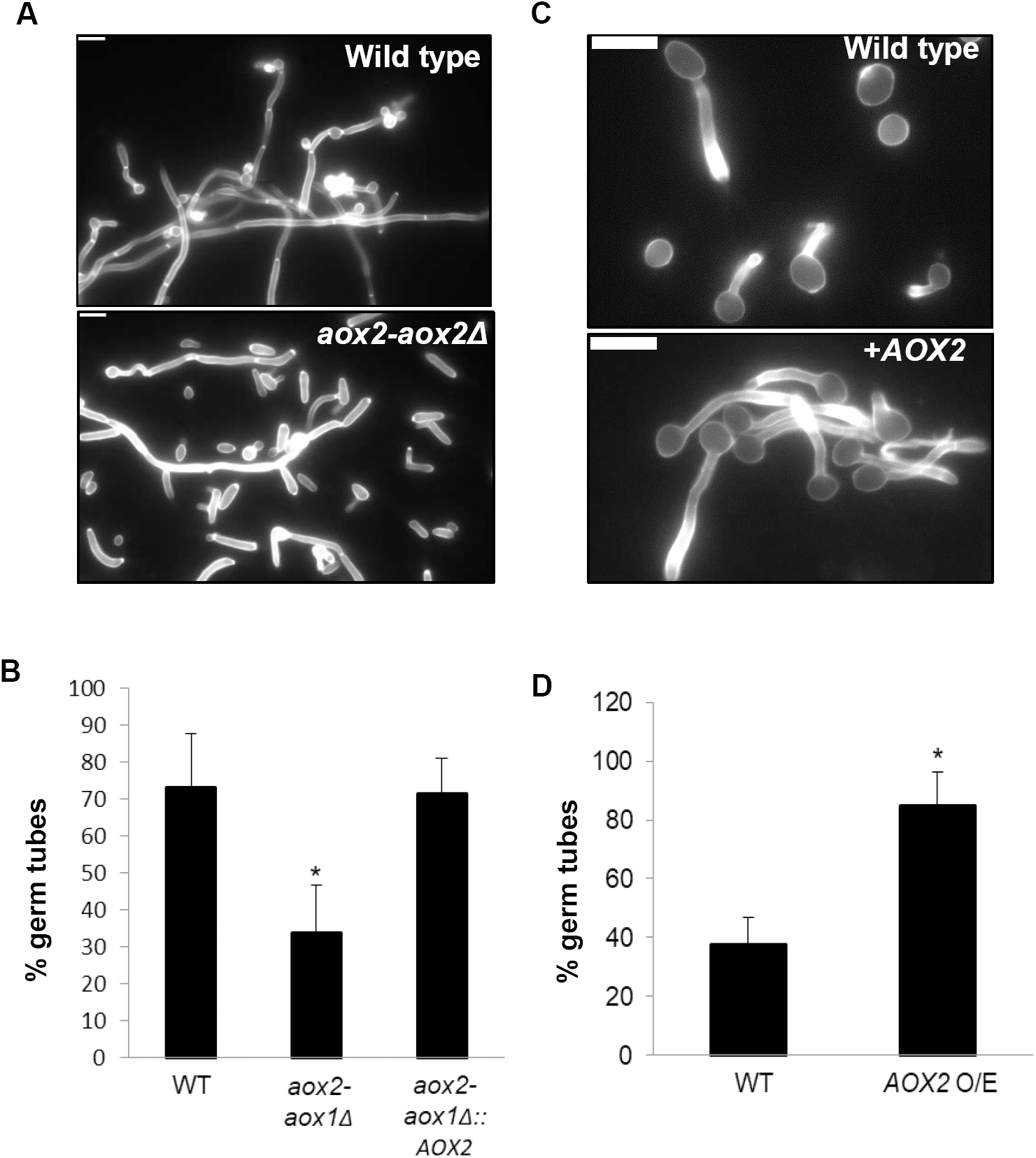
Aox2 levels affect the yeast-to-hypha transition. **A.** Representative examples of Wild-type, *aox2-aox1*Δ mutant and *AOX2* re-integration strain overexpressing *AOX2* were grown in RPMI at 37 °C for 16 h and **B**, analysis of percentage of hyphal cells from three independent experiments. **C.** Representative examples of wild-type and *AOX2* overexpression strains grown in serum-containing media for 90 min and **D,** analysis of percentage of hyphal cells from three independent experiments. At least 500 cells were counted per experiment. Results are presented as mean ± standard deviation. Student’s t-test was used to compare groups, *p<0.01.

### Aox is involved in the regulation of genes required for filamentous growth and glucose transport

As we observed that loss of *AOX2* could lead to profound effects under conditions of normal ETC function we examined the effects of its deletion upon global transcription during normal growth in YPD at 30 °C. For comparison we also examined the effects of addition of the Aox inhibitor SHAM to wild type cells. Interestingly the deletion of *AOX2* alone led to the differential expression of a number of genes under normal growth conditions (Figure 5A and 5B, Supplementary table S3). A number of genes affecting filamentous growth were differentially expressed in *aox2*Δ cells when compared to wild type, supporting our observation that Aox plays a role in hyphal transition. Genes involved in the repression of filamentous growth or genes which, when deleted, have hyperfilamentous phenotypes, including *CDC7, SKO1, RFG1, GAL10, PDE2* and *FGR22*, were upregulated (Alonso-Monge et al., 2010; Jung et al., 2005; Kadosh and Johnson, 2001; Lai et al., 2016; Singh et al., 2007; Uhl et al., 2003), while *DFG5*, encoding a GPI-anchored cell wall protein which promotes hyphal growth (Spreghini et al., 2003), was downregulated.

**Figure 5.**
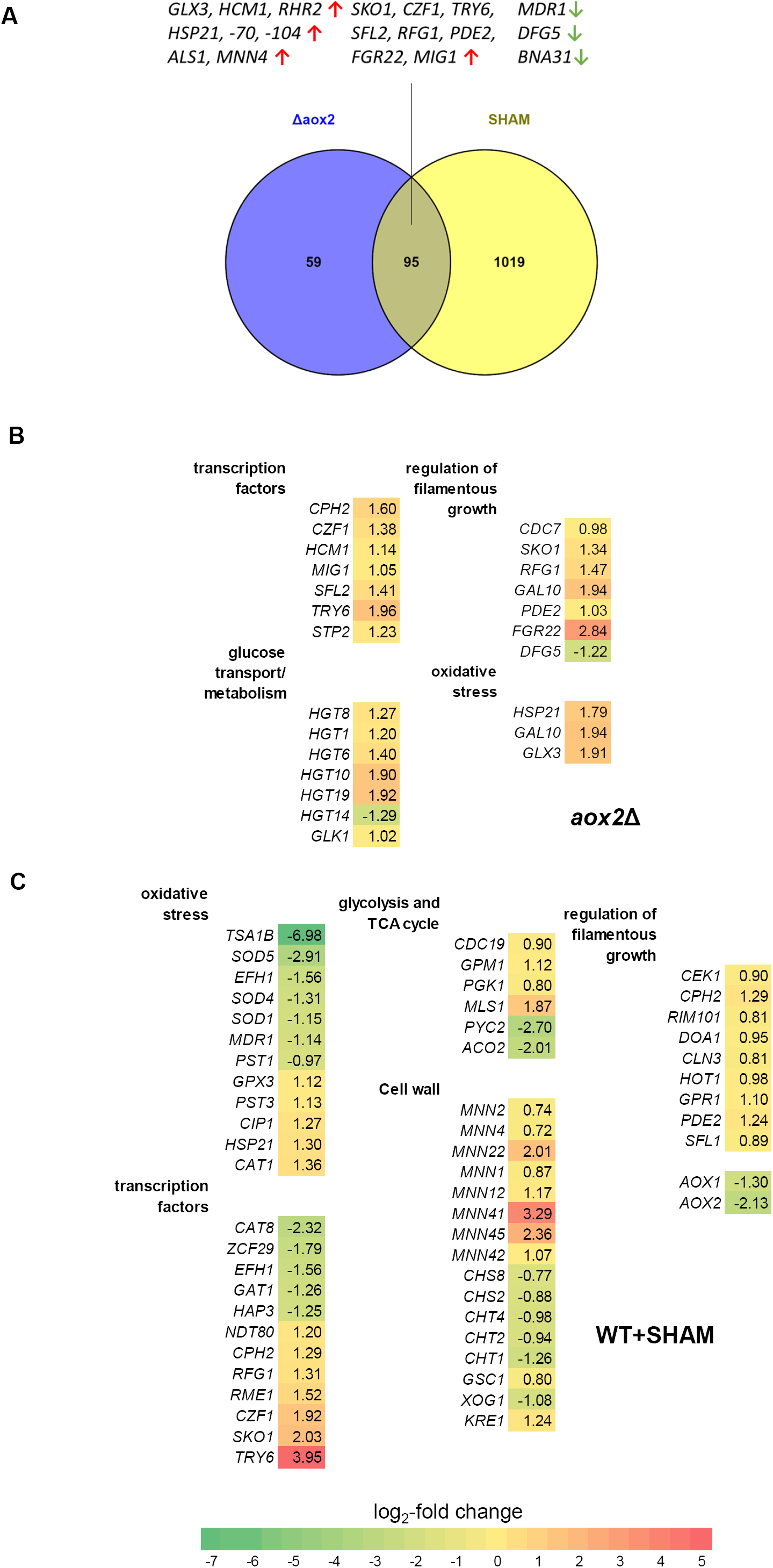
Global transcription changes upon deletion of AOX2 or treatment with SHAM. **A.** Overlap in differentially expressed genes between *aox2*Δ and SHAM-treated cells by RNAseq analysis. A selection of differentially expressed genes in **B**, SHAM-treated cells and **C**, *aox2*Δ. Genes were grouped by GO term, with log_2_-fold change vs. untreated wild-type shown. Green depicts downregulated genes and red/yellow depicts upregulated genes. Differentially expressed genes were identified by Cuffdiff v2.1.1, with q<0.05 being considered statistically significant.

Several members of the high-affinity glucose transporter gene family (Fan et al., 2002) were upregulated upon *AOX2* deletion (*HGT1, -6, -8, 10, 19*), suggesting increased glucose assimilation. This may be accompanied by an increase in glycolysis as the glucokinase gene *GLK1* was also upregulated. Interestingly, all three genes involved in the degradation of galactose (Leloir pathway) were upregulated (*GAL10, GAL1, GAL7)*. *GAL10* has been shown to have a role in regulating morphogenesis even in the absence of galactose, and the *GAL10* null mutant is hyperfilamentous (Singh et al., 2007). Upregulated genes associated with increased resistance to oxidative stress were also upregulated including *GLX3, GAL10 and HSP21* (Figure 5B).

SHAM treatment induced more gene expression changes than deletion of *AOX2*, suggesting additional off-target effects (Figure 5A, Supplementary Table S4). Upregulated transcription factors common to both *aox2*Δ and SHAM-treated groups included *CZF1, SFL2, SKO1* and *RFG1* which all regulate aspects of hyphal growth. *MDR1*, encoding the multidrug efflux pump, was downregulated approximately twofold in both *aox2*Δ and SHAM-treated groups, suggesting alternative respiration might play an important role in the response to antifungals. Transcripts of heat-shock factors Hsp21, -70, -104, which have been shown to have roles in virulence and stress responses other than heat-shock (Fiori et al., 2012; Mayer et al., 2012), were also upregulated in both groups.

A number of genes with functions in mannan biosynthesis and organisation were upregulated in SHAM treated cells (Figure 4C). Genes involved in β-glucan biosynthesis (*GSC1, KRE1*) were also upregulated. In contrast, a number of genes involved in chitin synthesis and degradation were downregulated. These transcriptional changes in cell wall genes observed with SHAM treatment could lead to a change in cell wall organisation. By comparison, there were fewer differentially expressed cell wall genes in the *aox2*Δ group, suggesting that off-target effects of SHAM, rather than inhibition of alternative respiration, may influence expression of cell wall genes.

*TRY6*, one of the major transcriptional activators of glycolysis genes, was the most highly upregulated transcription factor in both groups. In the SHAM-treated group, several glycolysis genes were upregulated (*CDC19, PGK1, GPM1*) as well as the glyoxylate cycle gene (*MLS1*), while some TCA cycle genes were downregulated (*PYC2, ACO1*). In the SHAM treated group, a number of genes involved in the oxidative stress response were differentially expressed, including downregulation of *TSA1B, SOD1, SOD4 and SOD5*. On the other hand, a number of oxidative stress genes were upregulated, including *CAT1* and *GPX3*. Differentially expressed genes linked to oxidative stress resistance in *aox2*Δ (*GLX3, GAL10 and HSP21*) may not necessarily be indicative of oxidative stress as their roles are indirect and they may have served other functions, in contrast to peroxidase and catalase genes in the SHAM group.

Analysis of the differentially expressed gene list from the SHAM treated group using PathoYeastract (Monteiro et al., 2017) showed that the transcription factor Ndt80 showed the most number of regulatory associations with the genes in the list based on documented DNA binding and expression evidence. *NDT80* was itself upregulated 2.2-fold. This transcription factor has a role in hyphal growth and biofilm formation (Lin et al., 2013; Sellam et al., 2010) and in sterol metabolism and drug resistance (Sellam et al., 2009). Other regulators that featured in this transcription factor ranking analysis include Tye7, which regulates glycolysis and carbohydrate metabolism, and Mcm1 and Sko1, which control hyphal growth (Alonso-Monge et al., 2010; Rottmann et al., 2003).

Similar to the *aox2*Δ results, a number of genes which negatively regulate hyphal growth were upregulated in the SHAM group (*DOA1, CLN3, PDE2 and SFL1*) (figure 4C) (Bauer and Wendland, 2007; Chapa et al., 2005; Jung et al., 2005; Kunze et al., 2007). These data suggest that SHAM treatment may suppress the yeast-to-hypha transition. In support of this we observed that SHAM was able to prevent hyphal transition on RPMI agar, a strong inducer of hyphal induction (Supplementary Figure 4). A similar effect of SHAM has been reported in other studies (Huh and Kang, 2001; Konno et al., 2006).

### Loss of Aox reduces virulence in zebrafish but not in the mouse model of systemic candidiasis

To examine the effect of the absence of alternative respiration on virulence, we employed the zebrafish model of systemic candidiasis. Zebrafish larvae were injected with 100 cfu or 500 cfu *C*. *albicans*. The fish were then kept at 28 °C and survival was assessed by inspection of heartbeat and motility after 24 and 48 hours. Fish in the *aox2-aox1*Δ group showed a higher survival rate than the wild-type strain (Figure 6A and 6B) but were not avirulent as all fish died after 48 h when 500 cfu *C*. *albicans* were injected (Figure 6B). A lower mortality rate was observed for both *aox2-aox1*Δ and *aox2*Δ deletion strains when compared to the wild-type after 24 hours (Figure 6A).

**Figure 6.**
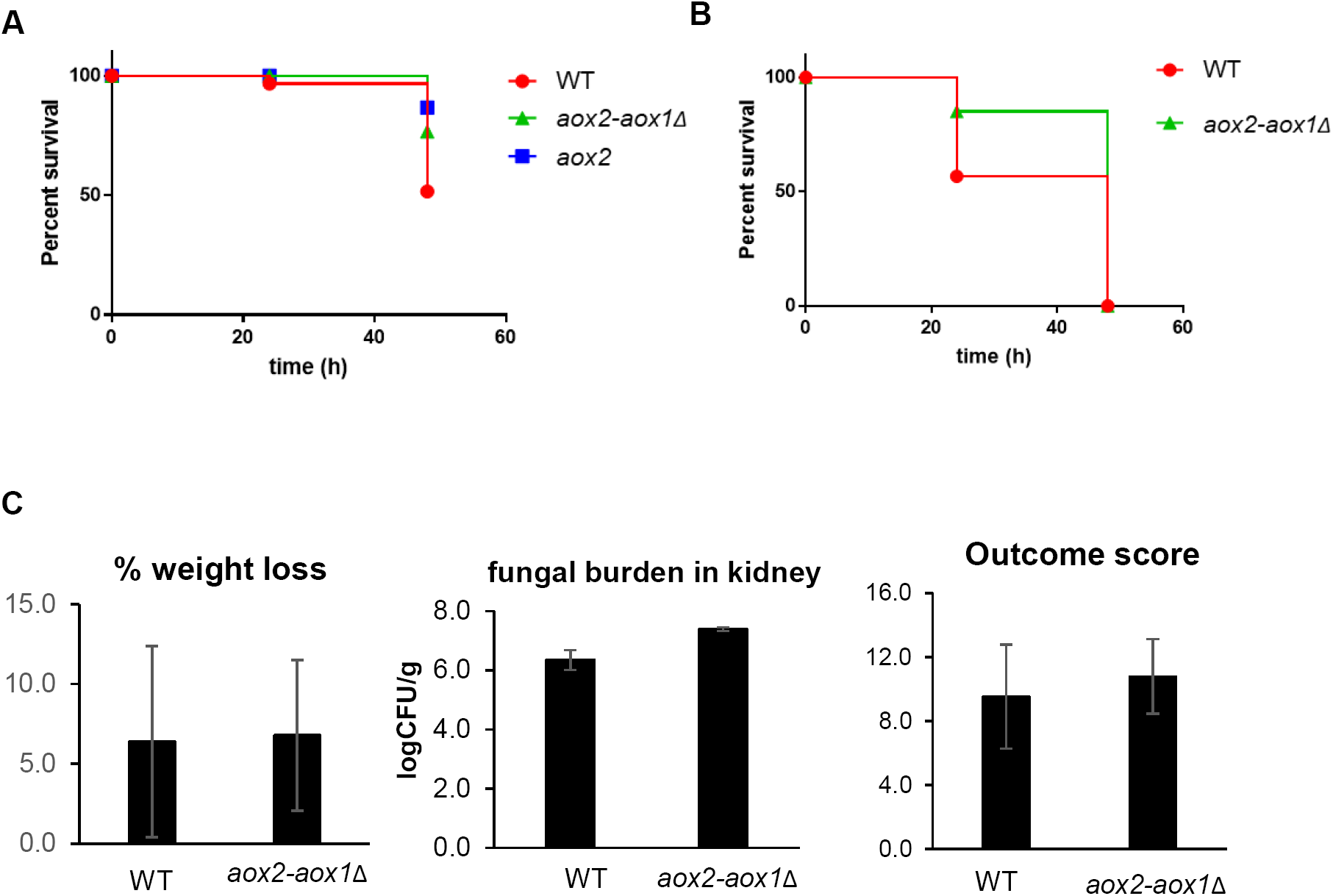
Deletion of *AOX2* causes decreased virulence in the zebrafish model of systemic candidiasis but not in the mouse model. Two day-old zebrafish larvae were injected with **A.** 100 cfu or **B.** 500 cfu *C*. *albicans* wild-type, *aox2*Δ or *aox2-aox1*Δ cells. Survival curves constructed from data collected over a 48 h period are shown. **B.** Wild-type or *aox2-aox1*Δ cells were used within a murine infection model as described in materials and methods. Percentage weight loss, fungal kidney burden and calculated outcome score are presented as mean ± standard deviation, n=6 mice per group. Student’s t-test was used to compare groups.

Based on the results of the zebrafish survival studies, we wished to determine the virulence of the *aox2-aox1*Δ mutant in the mouse model of systemic candidiasis. Mice were injected with *aox2-aox1*Δ via the tail vein and weight loss was monitored over the course of infection. There was no significant difference in weight loss between groups injected with *aox2-aox1*Δ and the wild-type strain (Figure 6C). The fungal burden in the kidneys was slightly higher in the *aox2-aox1*Δ group but this was not statistically significant (Figure 6C). The combination of these factors resulted in no significant difference in outcome score (MacCallum et al., 2010) between the *aox2-aox1*Δ and wild type groups. Therefore, the alternative pathway seems to be dispensable for virulence in the mouse model under the conditions tested.

### Pyocyanin inhibits alternative respiration and is synergistic with cyanide to inhibit respiration in *C*. *albicans*

Our data suggests that Aox may be important for cellular responses to respiratory challenge. It may be the case, therefore, that Aox function is important in the interaction between *C*. *albicans* and microbes that secrete factors that target respiratory machinery. An example of such an interaction lies between *C*. *albicans* and the pathogen *P*. aeruginosa, which produces cyanide under conditions of hypoxia and at high levels within cystic fibrosis patients (Cody et al., 2009). Alternative respiration may be required in this case as both species often occupy the same niche in cystic fibrosis patients (Chen et al., 2014). *P*. *aeruginosa* cells also produce molecules of the phenazine class and one of these, Pyocyanin (PYO), been shown to inhibit *C*. *albicans* filamentation (Morales et al., 2013).

We hypothesised that PYO might interfere with alternative respiration in synergy with cyanide to inhibit *C*. *albicans* respiration and halt growth of the yeast. We therefore tested the effects of PYO on alternative respiration following exposure to cyanide (Figure 7A and B). Two experiments were conducted in parallel: in the first, KCN was added to induce alternative respiration, at which point PYO was added to examine its effects on alternative pathway inhibition (Figure 7A). In the second experiment, PYO was added first, followed by KCN, to determine whether PYO alone affected respiration and also examine its effects on Aox2 induction (Figure 7B). The addition of PYO did not have any effect on routine respiration level, but significantly reduced the induction of Aox activity (Figure 7B). Furthermore, PYO was able to inhibit alternative respiration once it was induced by KCN (Figure 7A and 7C). These results show that PYO can act to inhibit of alternative respiration and also suggest that it may act in combination with cyanide to inhibit respiration in *C*. *albicans*.

**Figure 7.**
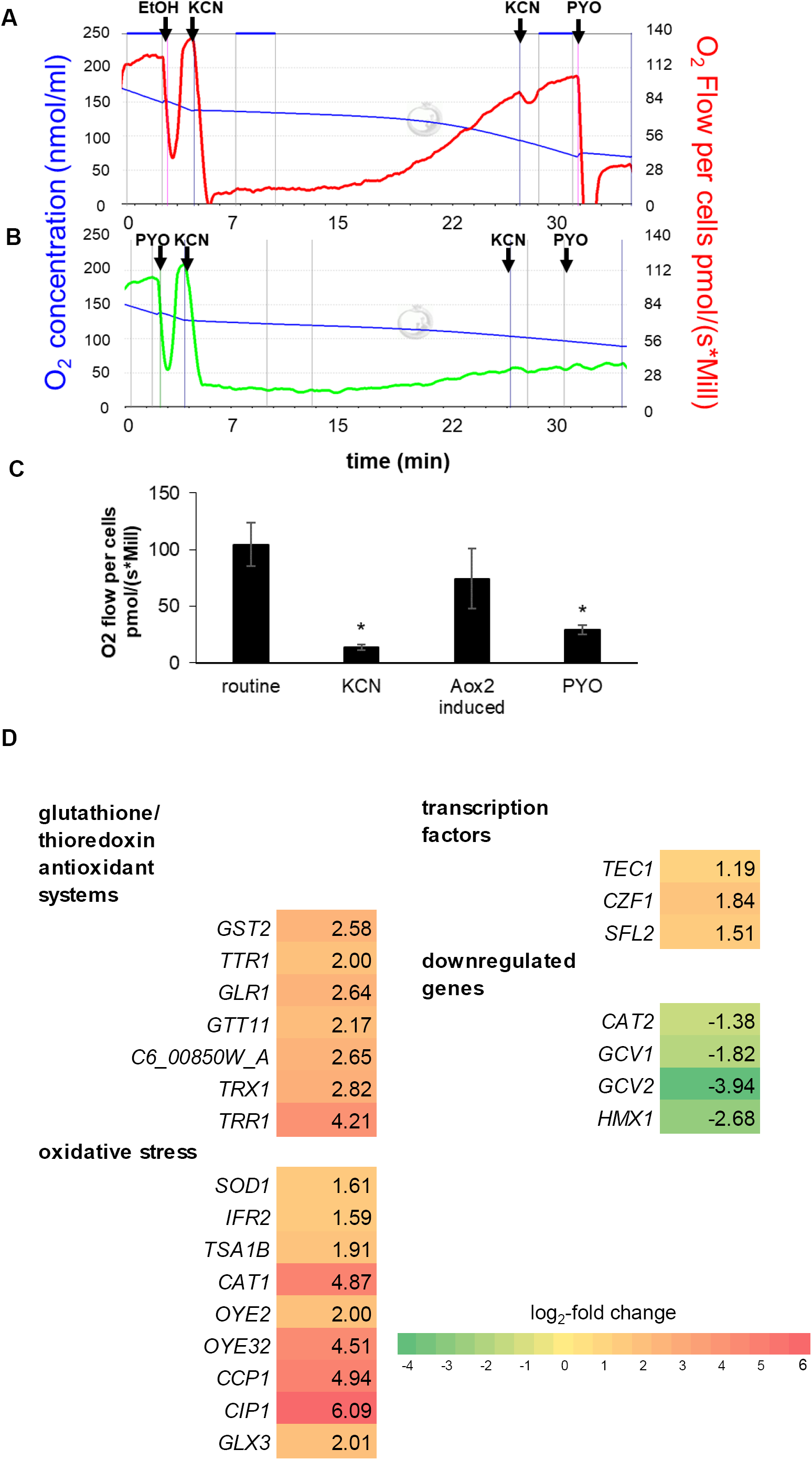
Pyocyanin inhibits alternative respiration. **A.** The effect of pyocyanin (PYO) on alternative respiration was investigated in wild-type *C*. *albicans*. Routine respiration was measured immediately before the addition of drugs. KCN was added to final concentration of 0.5 mM and KCN-inhibited respiration was recorded after 5 min. KCN was added a second time as indicated to a final concentration of 1 mM to confirm cyanide-insensitive respiration, after which Aox2-induced respiration was recorded. PYO was added to final concentration of 80 µM, after which PYO-inhibited respiration was measured. **B.** The effect of PYO on induction of Aox activity was investigated concurrently using the same schedule of drug addition and measurements as in **(A)**, except that PYO was added initially as indicated to a final concentration of 80 uM (an equal volume of the solvent ethanol was added in **(A)** as a control). The blue line shows oxygen concentration and red/green lines show the O_2-_flow per cells. **C.** Summary of results from four independent respirometry experiments performed as in **(A)**. Results are presented as mean ± standard deviation, Student’s t-test was used to compare groups, *p<0.05. **D.** A selection of differentially expressed genes in cells treated with Pyocyanin (80µM) for 30 min when compared to untreated is presented. Genes are grouped by GO term, with log_2_-fold change vs. untreated wild-type shown. Green depicts downregulated genes and red/yellow depicts upregulated genes. Differentially expressed genes were identified by Cuffdiff v2.1.1, with q<0.05 being considered statistically significant.

We next performed RNAseq to investigate the transcriptional changes made in *C*. *albicans* in response to PYO treatment. A small number of changes were observed upon a 30 min treatment with PYO. Genes involved in the glutathione- and thioredoxin antioxidant systems, as well as other oxidative stress response genes were upregulated, suggesting that PYO treatment causes oxidative stress in *C*. *albicans* (Figure 7D). Downregulated genes in PYO-treated cells include those involved in glycine catabolism (*GCV1* and *GCV2*), intracellular acetyl-CoA transport (*CAT2*), and utilisation of hemin iron *(HMX1*). Genes that were significantly differentially expressed in PYO treated cells are listed in Supplementary Table S5.

## Discussion

Although Aox function has been well studied in some filamentous fungi (Magnani et al., 2007; Nargang et al., 2012; Scheckhuber et al., 2011), the roles, regulation and contribution of alternative respiration in yeasts remains poorly understood. We present evidence that Aox has a role in protection of respiratory function, morphogenesis and in stress responses that may facilitate *C*. *albicans* in its capacity as both a commensal organism and as an opportunistic pathogen.

*C. albicans* Aox function would appear to be particularly important upon exposure to chemicals that disrupt ETC function. It may be the case that this reflects regular exposure to such compounds via interactions with host cells and other microbes. An example of such a compound is nitric oxide (NO). The inducible nitric oxide synthase (iNOS or NOS2) is the major source of NO that is generated by phagocytes as an antimicrobial defence. Interestingly data from *NOS2* knockout mice suggests that this enzyme is dispensible for clearance of oral candidiasis and for killing by macrophages (Farah et al., 2009). In addition nitrosative stress response genes were not found to be upregulated in *C*. *albicans* isolated from infected mice (Thewes et al., 2007). Deletion of *YHB1*, a nitric oxide dioxygenase involved in NO detoxification, from *C*. *albicans* led to a reduction in virulence in a mouse model but this loss did not correlate with *NOS2* function, also suggesting that NO production does not have a major effect on virulence in this model of candidiasis (Hromatka et al., 2005). In contrast, it was shown that *C*. *albicans* secretes factors specifically to inhibit NO production by immune cells (Collette et al., 2014). Why *C*. *albicans* might suppress NO production when it does not seem to be an important factor in determining virulence in a bloodstream infection is unclear, and highlights a lack of understanding of the precise roles of host-derived NO against *C*. *albicans*. The virulence of cells devoid of AOX function was not affected in a mouse infection model in our studies, also suggesting that the production of host NO is not a determinant of virulence in this model, or at least that the purpose of NO is not to inhibit *C*. *albicans* respiration. One possibility is that instead of inhibiting respiration, the fungicidal effects of NO may be derived from the generation of reactive nitrogen species such as peroxynitrite which forms when NO reacts with superoxide. NO-producing macrophages kill *C*. *albicans* more effectively when stimulated to produce higher levels of superoxide, suggesting that NO alone is not candidacidal (Vazquez-Torres et al., 1996).

Nevertheless our data clearly demonstrates that NO exposure leads to the inhibition of ETC function and to the upregulation of Aox which in turn facilitates continued respiration. One possibility is that NO exposure within the host elicits an Aox dependent stress response that primes *C*. *albicans* cells to initiate hyphal growth. In support of this idea, the contribution of the alternative pathway to total respiration in hyphal cells was shown to be significantly higher than in yeast (Guedouari et al., 2014). In addition, an inducer of Aox activity, guanosine-5’-monophosphate (GMP) (Milani et al., 2001), was shown to trigger filamentation in *C*. *albicans* under conditions of pH 4 in RPMI medium, in which hyphal growth is normally suppressed (Konno et al., 2006). It may be the case that Aox support of respiration allows TCA cycle to function under conditions of ETC inhibition and that this is important in the support of hyphal switching or hyphal growth. Communication between respiration and the TCA cycle seems to be a crucial aspect of hyphal growth, as inhibition of complex II to transfer electrons to ubiquinone using Thenoyltrifluoroacetone (TTFA) was shown to completely block filamentation (Watanabe et al., 2006). Further support comes from our finding that NO induced Aox expression, and indeed the overexpression of Aox in the absence of NO, promotes filamentation (Duvenage et al., 2018). The RNAseq data presented here also supports the hypothesis that AOX has a role in hyphal switching, as several genes involved in its regulation were differentially expressed in both the *aox2*Δ and SHAM treated groups. For example, the transcription factor *SKO1*, upregulated in both groups, represses the hyphal transition by controlling the expression of hyphal-specific genes such as *HWP1* and *ECE1* (Alonso-Monge et al., 2010).

As Aox appears to play an important role in mediating filamentation we would expect that the loss of Aox, or indeed its inhibition, would have a marked effect on *C*. *albicans* virulence. Our experimental approach tested this hypothesis in both zebrafish and mouse models of infection. The zebrafish larva survival model focuses on the innate immune response to *C*. *albicans*, as adaptive immunity has not developed at the time of infection. In this case infection with either *aox2*Δ or *aox2-aox1*Δ mutant strains showed attenuated virulence. However, the fact that no larvae survived longer than 48 h – in the case where a higher number of CFUs were injected – showed that the loss of alternative respiration merely slowed the progress of the infection. This could be due to a slower emergence of hyphae at the incubation temperature of 28 °C. A correlation between filamentation and increased virulence in this model has been noted previously (Mallick et al., 2016).

Another reason for the maintenance of Aox function may reside with interactions with other microbes that secrete molecules that target classical respiration. Such interactions may form part of symbiotic or competitive interactions between resident or invading microorganisms. One such example is the interaction between *C*. *albicans and P*. *aeruginosa*, which are frequently co-isolated from the cystic fibrosis lung. In cystic fibrosis *P*. *aeuroginsa* is known to secrete cyanide, as well a class of molecules known as phenazines – including pyocyanin (PYO) – which inhibits filamentation in *C*. *albicans* (Morales et al., 2013). We chose to examine the effect of the phenazine pyocyanin (PYO) on respiration in *C*. *albicans* based on the observed sensitivity of the *aox2-aox1*Δ mutant to its thioanalogue, methylene blue (MB). Although both PYO and MB have pleiotropic effects, current evidence agrees that one of their targets is the ETC (Kasozi et al., 2011; Schirmer et al., 2011). In our RNAseq analysis of PYO treated cells the most noticeable trend was an oxidative stress response, with upregulation of genes in the glutathione and thioredoxin antioxidant systems. Induction of oxidative stress and disruption of the glutathione redox cycle by pyocyanin has previously been reported in human endothelial cells (Muller, 2002). As one of the targets of PYO is mitochondria, it is also possible that mitochondrial dysfunction could have contributed to increased oxidative stress.

The production of cyanide by *P. aeruginosa* may be another strategy to inhibit growth of competing microorganisms such as *C*. *albicans*, however alternative respiration allows for normal growth in the presence of cyanide. Our data suggests that mitochondria respiring via the alternative pathway are more susceptible to the effects of PYO. The production of a second factor, PYO, by *P*. *aeruginosa* to counteract alternative respiration in synergy with cyanide, highlights the relevance of *C*. *albicans* Aox function in microbial interactions where electron transport chain inhibition can occur. The function of Aox in resistance to respiratory stress is also likely one of its physiological functions in its co-existence with competing microbes such as *P*. *aeruginosa*. Whether elevated Aox activity influences filamentation in these situations should be considered in future work.

In summary we find that Aox plays an important role in the responses of *C*. *albicans* to ETC inhibition. The induction of Aox in support of alternative respiration is clearly essential for growth and survival as *C*. *albicans* depends upon mitochondrial function for growth. Our findings suggest that, as has been found in other eukaryotic organisms that maintain Aox function, alternative respiration is likely to facilitate survival and adaptability within specialised niche environments that commonly damage ETC function.

## Supplementary figures

**Figure S1.**
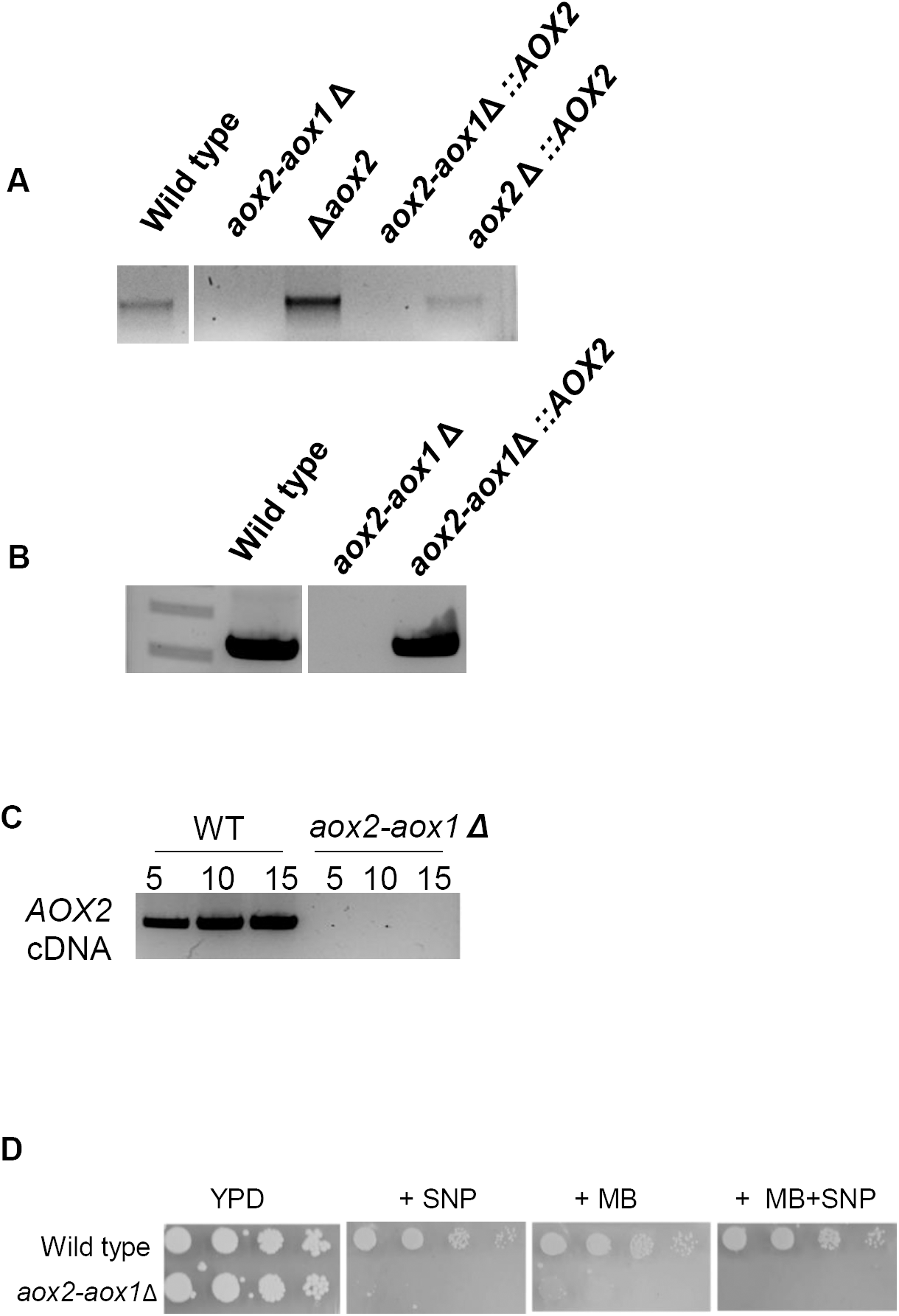
Confirmation of deletion of *AOX* genes. Deletion of *AOX2* and *AOX2-AOX1* region. **A.** PCR to confirm the disruption of *AOX1* in the *aox2-aox1*Δ mutant and corresponding *AOX2* overexpression strain, using primers AOX1 flank F and AOX1 flank R. **B.** PCR to confirm disruption of *AOX2* in the *aox2-aox1*Δ mutant and reintegration in the corresponding *AOX2* overexpression strain, using primers AOX2 ORF F and AOX2 ORF R**. C.** RT-PCR for the presence of *AOX2* transcripts in the wild-type strain and in *aox2-aox1*Δ from cells treated with 1 mM KCN for 1 hour, using increasing amounts of template cDNA. **D.** The wild-type or *aox2-aox1*Δ mutant were spotted onto plates containing 1 mM SNP or 10 µg/ml methylene blue. Growth was evaluated after 48 h. Results are representative of at least three independent experiments.

**Figure S2.**
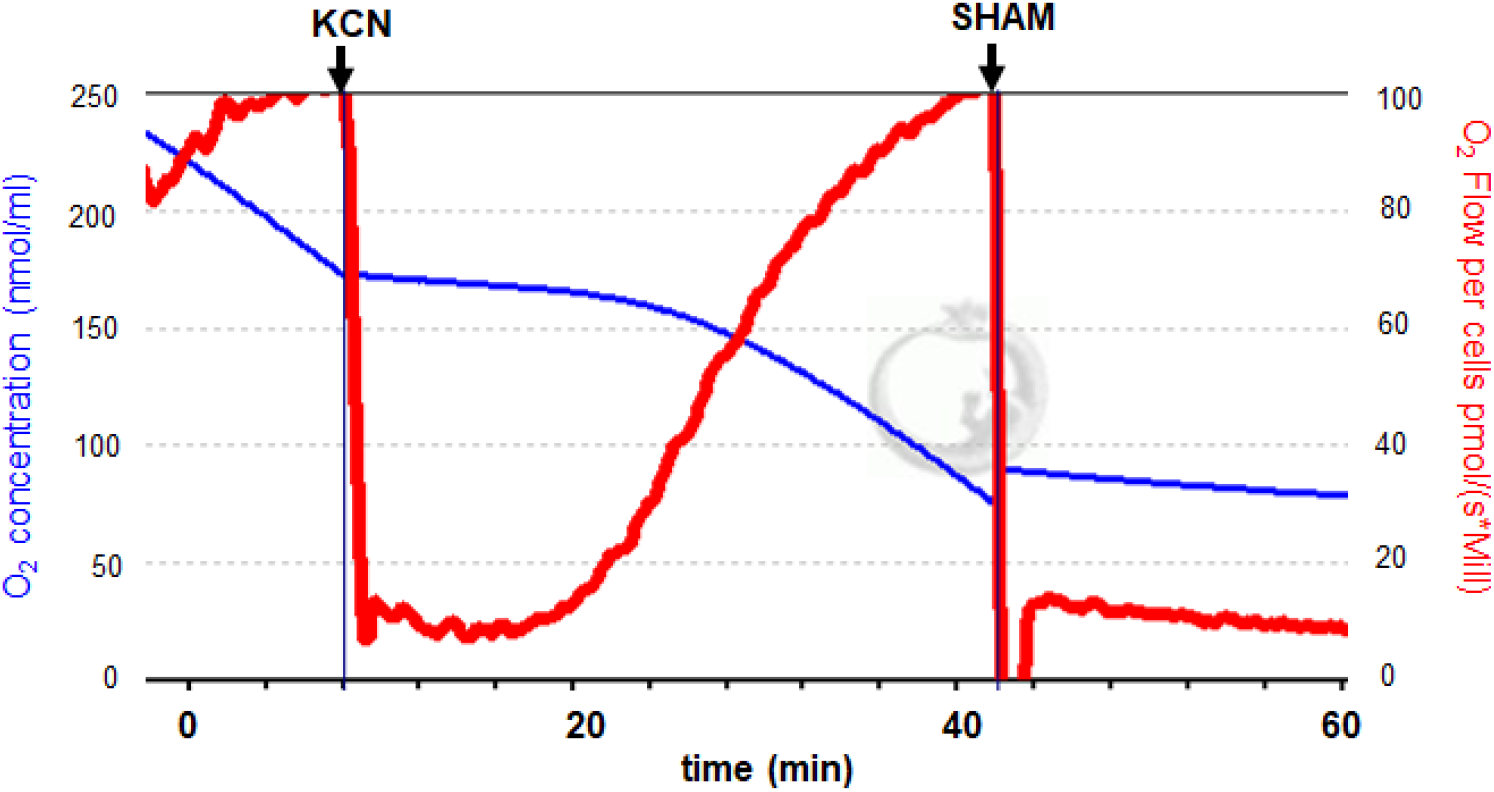
Whole cell respirometry data showing Aox induction after KCN inhibition. Inhibition of respiration with 1 mM KCN induces Aox activity within 20 minutes. Aox activity is inhibited by addition of 1 mM SHAM.

**Figure S3.**
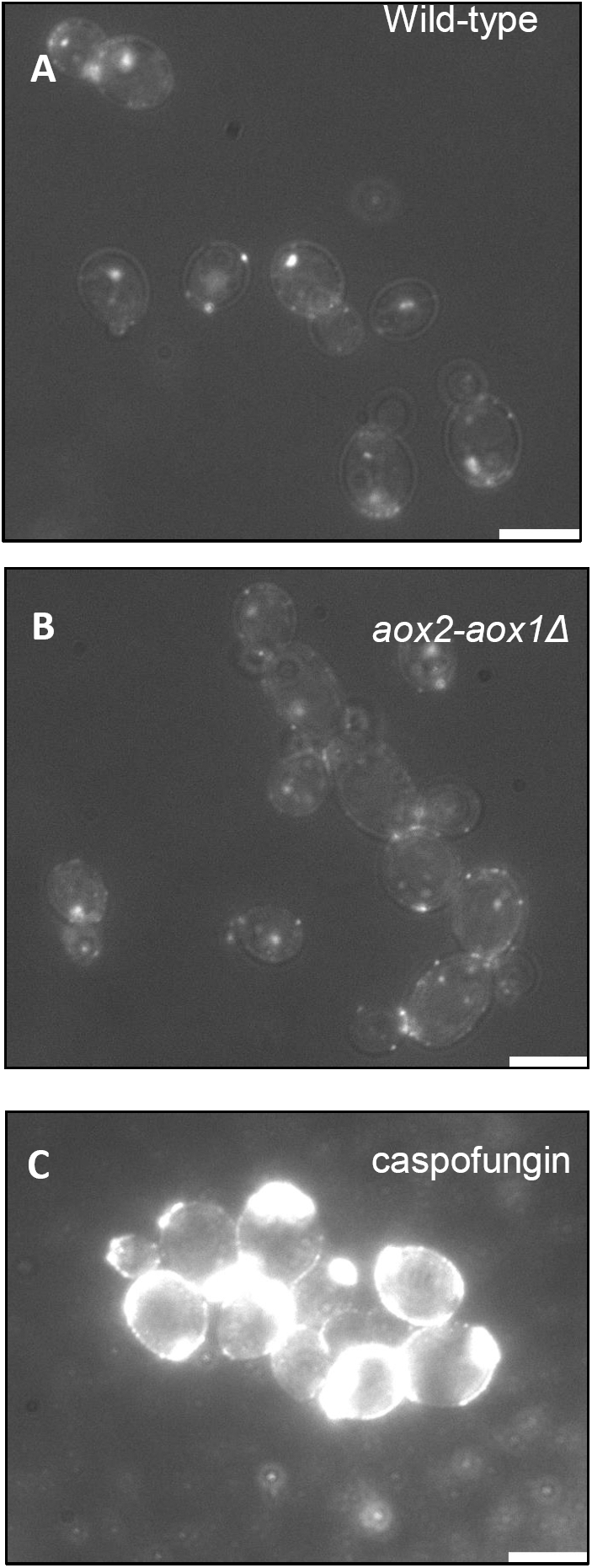
Dectin-1 staining of wild type and *aox2-aox1*Δ cells. Representative examples showing dectin-1 staining of **A.** wild-type SC5314 and **B.** *aox2-aox1*Δ *C*. *albicans*. Images were captured with GFP settings with low brightfield illumination. **C.** Wild-type *C*. *albicans* grown overnight in the presence of 10 ng/ml caspofungin overnight is shown as a positive control.

**Figure S4.**
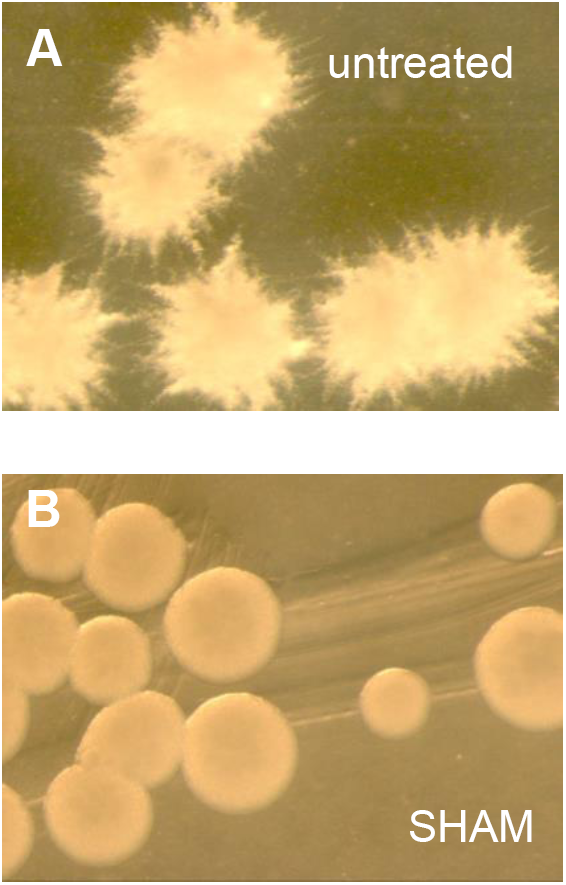
SHAM inhibits filamentation. Inhibition of filamentation by 3.2 mM SHAM during growth on RPMI agar at 37 °C for 48 h.

**Supplementary Table S1. *C. albicans* strains used in this study**

**Supplementary Table S2. Primers used in this study**

**Supplementary Table S3. Differentially expressed genes between wild-type and *aox2*Δ strains analysed by RNAseq**

**Supplementary Table S4. Differentially expressed genes between untreated SHAM-treated *C***. ***albicans* analysed by RNAseq**

**Supplementary Table S5. Differentially expressed genes between untreated pyocyanin-treated *C***. ***albicans* analysed by RNAseq**

**Supplementary data Table S1.** *C. albicans* strains used in this study

| Strain | Genotype | Source |
| --- | --- | --- |
| SC5314 | Typed strain | Typed strain |
| SN87 | *his/his1 leu2/leu2*<br>*URA3/ura3::imm434 iro1::IRO1/iro1::imm434* | [17] |
| SN152 | *arg4/arg4 his/his1 leu2/leu2*<br>*URA3/ura3::imm434 iro1::IRO1/iro1::imm434* | [17] |
| *aox2*Δ | *arg4/arg4 his/his1 leu2/leu2*<br>*URA3/ura3::imm434 iro1::IRO1/iro1::imm434*<br>*aox2::LEU2/aox2::HIS1* | This work |
| *aox2-aox1*Δ | *his/his1 leu2/leu2*<br>*URA3/ura3::imm434 iro1::IRO1/iro1::imm434 aox2-aox1::LEU2/aox2-aox1::HIS1* | This work |
| *aox2-aox1*Δ::AOX2 | As for Δ*aox2-aox1*, ADH1/*adh1::pNIM-AOX2* | This work |
| AOX2 overexpression | ADH1/*adh1::pNIM-AOX2* | This work |

**Supplementary data Table S2.**
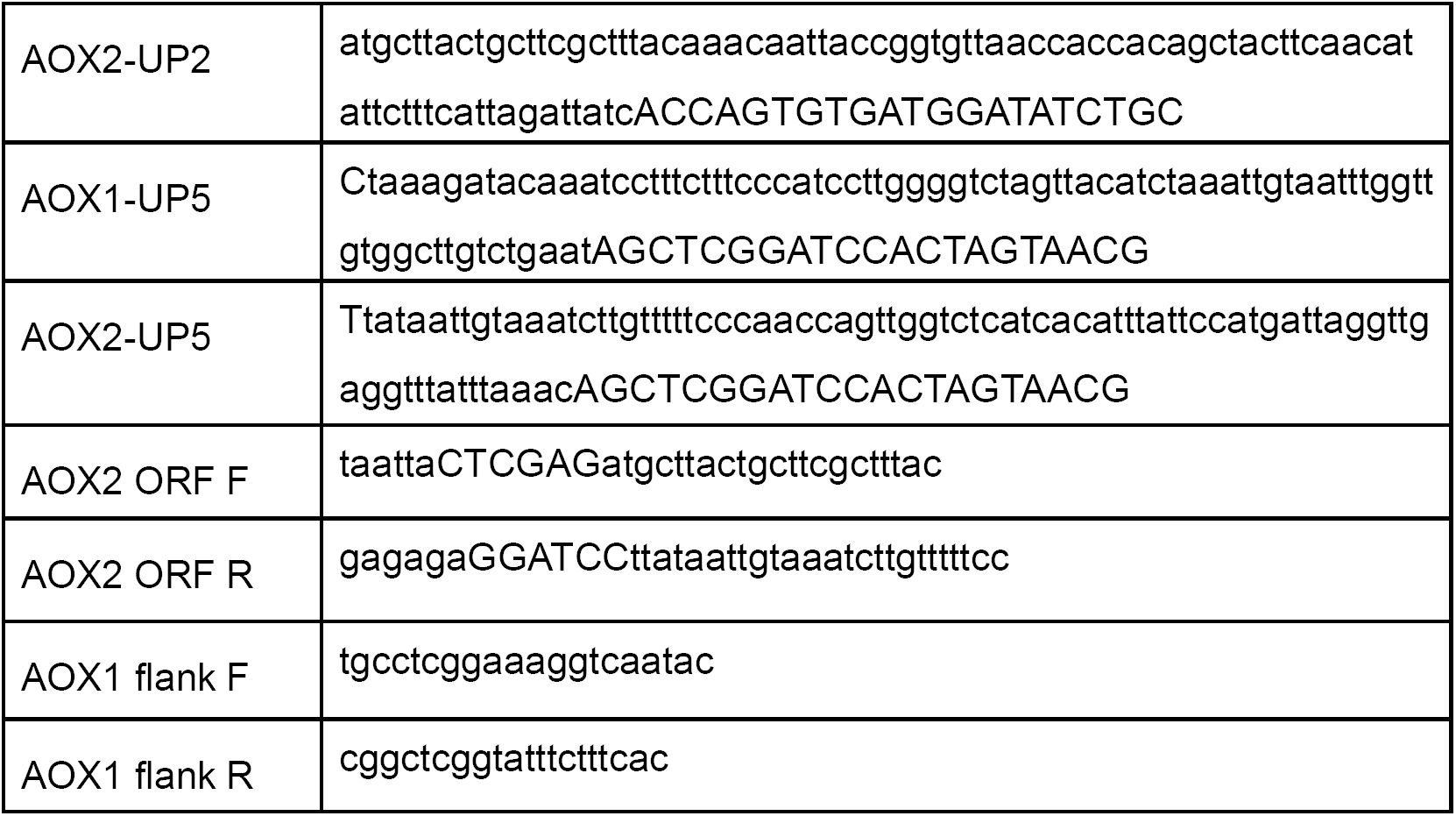
Primers used in this study

